# Behavioral effects of zonisamide on L-DOPA-induced dyskinesia in Parkinson’s disease model mice

**DOI:** 10.1101/2023.04.26.538384

**Authors:** Hiromi Sano, Atsushi Nambu

## Abstract

Zonisamide (ZNS; 1,2-benzisoxazole-3-methanesulfonamide) was initially developed and is commonly used as an anticonvulsant drug. However, it has also shown beneficial effects on Parkinson’s disease (PD), a progressive neurodegenerative disorder caused by the loss of dopaminergic neurons in the midbrain. Recent clinical studies have suggested that ZNS can also have beneficial effects on L-DOPA-induced dyskinesia (LID), which is a major side effects of long-term L-DOPA treatments for PD. In the present study, we examined the behavioral effects of ZNS on LID in PD model mice. Acute ZNS treatment did not have any observable behavioral effects on LID. Contrastingly, chronic ZNS treatment with L-DOPA delayed the peak of LID and reduced the severity of LID before the peak, but increased the duration of LID in a dose-dependent manner of ZNS, compared to PD model mice treated with L-DOPA alone. Thus, ZNS appears to have both beneficial and adverse effects on LID.

## Introduction

Parkinson’s disease (PD) is a neurodegenerative disorder caused by the loss of nigrostriatal dopaminergic neurons in the substantia nigra pars compacta (SNc), leading to motor and non-motor symptoms, including akinesia, bradykinesia, resting tremor, rigidity, cognitive dysfunctions, depression, and insomnia [1, 2]. Supplementation therapy with L-DOPA can effectively control symptoms in the early stage of PD (a well-controlled “honeymoon” period) [2], but long-term administration can induce wearing-off, a reduction of the effective time, and involuntary movements known as L-DOPA-induced dyskinesia (LID). Once LID is established, it is difficult to relieve involuntary movements without compromising antiparkinsonian effects [3]. The reduction and prevention of LID are major issues in PD drug therapy.

Zonisamide (ZNS; 1,2-benzisoxazole-3-methanesulfonamide) was initially approved for epilepsy treatment in Japan in 1989 and later approved worldwide [4]. It was also serendipitously discovered to have beneficial effects on the symptoms of PD [5-7]. When ZNS was administered to a patient with PD who incidentally had convulsive attacks, the attacks disappeared, and the symptoms of PD also improved. Several clinical trials have explored the effects of ZNS on the motor symptoms of PD, and in 2009, it was approved as an adjunctive treatment for patients with PD who responded insufficiently to L-DOPA treatment in Japan. ZNS significantly reduces the duration of wearing-off without worsening dyskinesia [8]. Although the mechanism of ZNS in the symptoms of PD has not been fully elucidated, it has been suggested to have multiple mechanisms of action, including inhibition of monoamine oxidase [9, 10], blockade of T-type calcium channels [11, 12], modulation of L-DOPA-dopamine metabolism [13], and neuroprotection [14, 15].

In our previous study, we investigated the behavioral and neurophysiological effects of ZNS on LID in PD model mice [16]. The chronic ZNS injection (100 mg/kg) in combination with L-DOPA increased the duration and severity of LID and altered neuronal activity in the output nuclei of the basal ganglia, i.e., the substantia nigra pars reticulata (SNr). This resulted in a decreased spontaneous firing rate, increased cortically evoked inhibition, and decreased cortically evoked late excitation, which may explain the mechanism of enhanced LID [16, 17]. In this study, we aimed to evaluate the behavioral effects of ZNS on LID in more detail by injecting L-DOPA or a combination of L-DOPA and ZNS at various doses to induce LID in PD model mice. We observed abnormal involuntary movements (AIMs) after L-DOPA with or without ZNS injection.

## Materials and Methods

### Animals

Male ICR mice were obtained from Japan SLC. All the mice used in the experiments were more than eight weeks old. The mice were maintained on a 12-hour light and dark cycle with *ad libitum* access to food and water. All experimental procedures were approved by the Institutional Animal Care and Use Committee of National Institutes of Natural Sciences.

### Experimental design

To investigate the effects of acute (Experiment 1, Fig 1A) and chronic (Experiment 2, Fig 1B) ZNS injection on LID, we performed the following experiments. First, mice received a 6-hydroxydopamine (6-OHDA) injection into the right medial forebrain bundle (MFB) to eliminate dopaminergic neurons. Two to three weeks later, motor deficits were evaluated using the cylinder test. In Experiment 1, after injecting L-DOPA for nine consecutive days, we observed dyskinetic behavior after L-DOPA with saline or ZNS injections (Fig 1A). In Experiment 2, after injecting L-DOPA with saline or ZNS for nine consecutive days, we observed dyskinetic behavior after L-DOPA with saline or ZNS injections (Fig 1B). One or two days after the behavioral test, the mice were perfused transcardially with formalin, and histological analysis was performed to evaluate the degeneration of dopaminergic neurons in the SNc.

**Fig 1.**
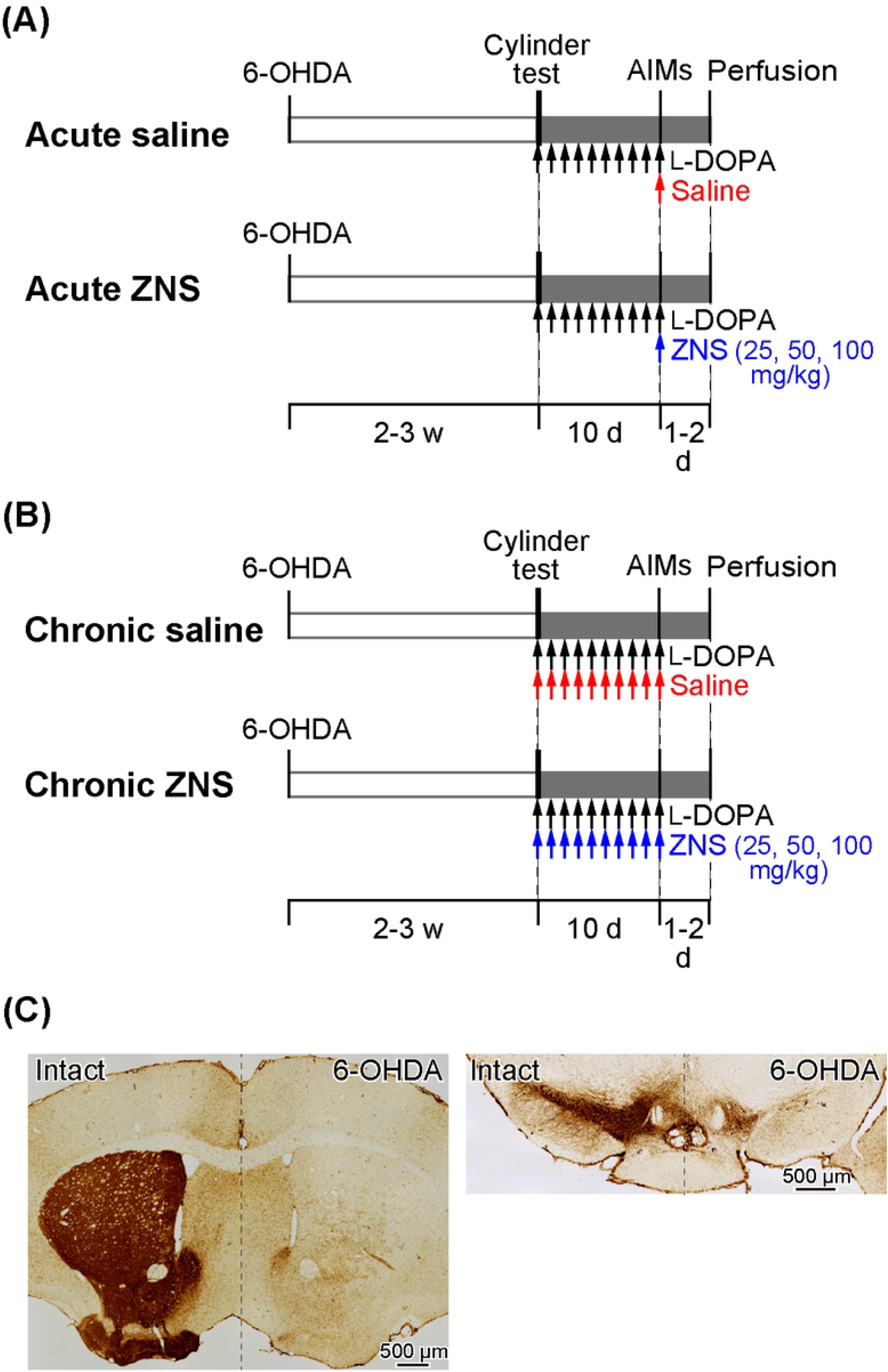
A schematic representation of the experimental design. (A and B) Timelines of Experiments 1 and 2. Mice received 6-OHDA infusions into the right medial forebrain bundle. The cylinder test was performed to evaluate motor deficits two to three weeks after injection. (A) Experiment 1 was performed to evaluate the acute effects of ZNS on LID. After the cylinder test, mice were injected with L-DOPA for nine consecutive days (20 mg/kg; black arrows), and behavioral tests for AIMs were conducted on the 10^th^ day after L-DOPA with saline (red arrow) or ZNS (25, 50, and 100 mg/kg; blue arrow) injections. (B) Experiment 2 was performed to evaluate the chronic effects of ZNS on LID. After the cylinder test, mice were injected with L-DOPA (20 mg/kg; black arrows) with saline (red arrows) or ZNS (25, 50, and 100 mg/kg; blue arrows) for nine consecutive days, and AIMs were observed on the 10^th^ day after L-DOPA with saline or ZNS injection. (C) Dopaminergic nerve terminals in the striatum (left panel) and dopaminergic neurons in the substantia nigra pars compacta (right panel) on the intact and 6-OHDA-lesioned sides were visualized with tyrosine hydroxylase immunohistochemistry. 6-OHDA, 6-hydroxydopamine; ZNS, zonisamide; AIMs, abnormal involuntary movements; LID, L-DOPA-induced dyskinesia

### 6-OHDA lesion

6-OHDA was infused as previously described [16, 17]. Briefly, desipramine hydrochloride (25 mg/kg, *i*.*p*.; Sigma-Aldrich) was injected into mice 30 min before the 6-OHDA infusion. Mice were anesthetized with isoflurane (1.0 – 1.5%) and fixed in a stereotaxic apparatus (SR-6M-HT, Narishige). The skin of the head was incised, and a small hole was made on the right side of the skull. We injected 6-OHDA-hydrobromide (4 mg/ml; Sigma-Aldrich) that was dissolved in saline containing 0.02% ascorbic acid into the right medial forebrain bundle at the coordinates 1.0-mm posterior, 1.4-mm lateral, and 4.8-mm ventral from the bregma, using a glass micropipette and a micro infusion pump (1 μl at 0.2 μl/min; Micro4, WPI).

### Cylinder test

To evaluate motor deficits induced by a unilateral lesion of the nigrostriatal dopaminergic neurons, a cylinder test was performed two to three weeks after the 6-OHDA infusion, as previously described (Figs 1A and 1B) [16, 17]. Each mouse was placed in a clear acrylic cylinder (10 cm in diameter and 20 cm in height) and was observed for 5 min without habituation to the cylinder. The number of contacts with the right or left forelimb was counted, and the limb-use asymmetry score was calculated as the number of wall contacts with the left forelimb (the forelimb contralateral to the lesion side) as a percentage of the total wall contacts. If mice did not show impairments in the usage of the left forelimb compared with the right forelimb, 6-OHDA was injected again so that the limb-use asymmetry score was close to 30%.

### L-DOPA and ZNS treatment

Daily L-DOPA treatment was initiated after the cylinder test. The 6-OHDA-lesioned mice were injected with L-DOPA (Experiment 1, Fig 1A) or L-DOPA with saline or ZNS (Experiment 2, Fig 1B) once for nine consecutive days. L-DOPA was injected (20 mg/kg, *i*.*p*.; Ohara Pharmaceutical Co., Ltd) in combination with benserazide (12 mg/kg, *i*.*p*.; Sigma-Aldrich). In Experiment 2, saline (10 ml/kg, *i*.*p*.) or different doses of ZNS (25, 50, and 100 mg/kg, *i*.*p*.; provided by Sumitomo Pharma Co., Ltd.) were injected 10 min after L-DOPA treatment.

### Behavioral test

On day 10, after injecting L-DOPA with or without ZNS treatments for nine consecutive days (Experiments 1 and 2, respectively), saline (10 ml/kg, *i*.*p*.) or different doses of ZNS (25, 50, and 100 mg/kg, *i*.*p*.) were injected 10 min after L-DOPA (20 mg/kg, *i*.*p*.) treatment. The mice were then placed in separate cages, and dyskinetic behaviors were observed every 20 min for 1 min over a 180-min period (Figs 1A and 1B). Dyskinetic behavior was scored based on the previously described AIMs scale [16, 17], which classifies AIMs into four subtypes: locomotive (increased locomotion with contralateral rotations), axial (contralateral dystonic posture of the neck and upper body towards the side contralateral to the lesion), limb (jerky and fluttering movements of the limb contralateral to the side of the lesion), and orolingual AIMs (vacuous jaw movements and tongue protrusions). Each of these for subtypes was scored on a severity scale ranging from 0 to 4 (0, absent; 1, occasional; 2, frequent; 3, continuous; 4, continuous, and not interruptible by external stimuli). Furthermore, we calculated the total AIMs score by adding up the four AIM subtypes.

### Histology

One to two days after the behavioral tests, the mice were deeply anesthetized with sodium thiopental (100 mg/kg, *i*.*p*.) and were perfused transcardially with 0.01M phosphate-buffered saline (PBS) followed by 10% formalin in 0.01M PBS. The brains were removed, postfixed in 10% formalin at 4 °C overnight, cryoprotected in 15% sucrose in 0.01M PBS at 4 °C, and then in 30% sucrose in 0.01M PBS at 4 °C. These brains were subsequently frozen and sectioned coronally into 40-μm sections. The free-floating sections were incubated with antibodies against tyrosine hydroxylase (TH; 1: 500; Millipore), and then visualized with biotinylated secondary antibodies and the ABC method [15].

### Statistical analysis

L-DOPA-induced AIMs were compared among the four groups (saline and different doses of ZNS) using a two-way analysis of variance (ANOVA) with repeated measures and Tukey’s post hoc test. Statistical analyses were performed using Prism7 software (GraphPad Software Inc.), and *p* < 0.05 was considered statistically significant.

## Results

TH-positive terminals in the striatum and TH-positive neurons in the SNc dramatically decreased on the 6-OHDA injection side, as shown in Fig 1C. In the cylinder test, mice treated with 6-OHDA showed a significant reduction in the usage of the forelimb contralateral to the lesion, and the limb-use asymmetry scores were decreased to 28.91 ± 6.09 % (mean ± S.D.), which was similar across the four groups in Experiments 1 and 2. These results confirmed that the dopaminergic neurons in the SNc were successfully lesioned, and that the mice exhibited the symptoms of PD.

### Effects of acute ZNS treatment on AIMs, Experiment 1

On day 10, after injecting L-DOPA for nine consecutive days, saline or ZNS (25, 50, and 100 mg/kg) was injected 10 min after L-DOPA treatment (Fig 1A), and AIMs were observed in different body parts, and were scored for 1 min at every 20 min after L-DOPA treatment (Fig 2). We compared the total AIMs scores between the acute saline and ZNS (25, 50, and 100 mg/kg) treatment groups. There was no statistically significant difference between all groups (Fig 2A; two-way repeated measures ANOVA: time, F (8, 280) = 169.4, *p* < 0.01; treatment, F (3, 35) = 0.36, *p* = 0.78; interaction, F (24, 280) = 1.83, *p* < 0.05). There was also no significant difference between all groups in the locomotive (Fig 2B; two-way repeated measures ANOVA: time, F (8, 280) = 99.68, *p* < 0.01; treatment, F (3, 35) = 0.65, *p* = 0.59; interaction, F (24, 280) = 1.81, *p* < 0.05), axial (Fig 2C; two-way repeated measures ANOVA: time, F (8, 280) = 114.1, *p* < 0.01; treatment, F (3, 35) = 0.38, *p* = 0.77; interaction, F (24, 280) = 1.08, *p* = 0.37), limb (Fig 2D; two-way repeated measures ANOVA: time, F (8, 280) = 82.34, *p* < 0.01; treatment, F (3, 35) = 1.64, *p* = 0.20; interaction, F (24, 280) = 1.80, *p* < 0.05), and orolingual (Fig 2E; two-way repeated measures ANOVA: time, F (8, 280) = 115.3, *p* < 0.01; treatment, F (3, 35) = 1.64, *p* = 0.20; interaction, F (24, 280) = 2.51, *p* < 0.01) AIMs scores. We compared the peaks of AIMs in the acute saline and ZNS (25, 50, and 100 mg/kg) treatment groups (inverted triangles in Fig 2). Peaks of the total, locomotive, axial, limb, and orolingual AIMs scores were distributed between 40 and 100 min. These results suggest that acute ZNS injections did not affect L-DOPA-induced AIMs in PD model mice.

**Fig 2.**
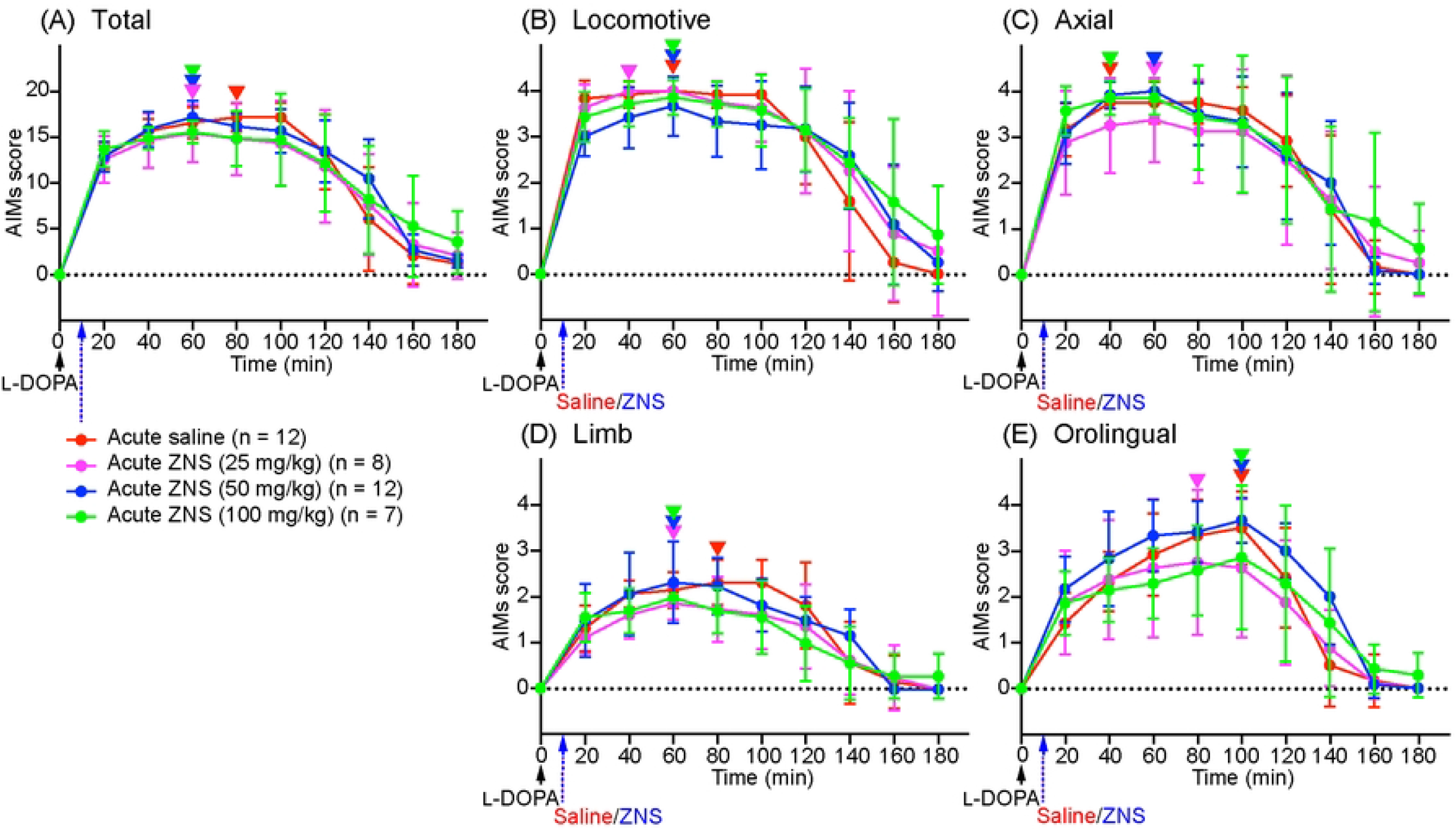
Acute ZNS treatment does not affect AIMs scores. Total (A), locomotive (B), axial (C), limb (D), and orolingual (E) AIMs were scored for 180 min at every 20 min after injecting L-DOPA (20 mg/kg; black arrow) at time 0 min with saline (red) or ZNS (25 mg/kg, magenta; 50 mg/kg, blue; 100 mg/kg, green) at 10 min. Data are expressed as mean ± SD. Peaks of AIMs with saline or ZNS are indicated by inverted triangles with corresponding colors. ZNS, zonisamide; AIMs, abnormal involuntary movements

### Effects of chronic ZNS treatment on AIMs, Experiment 2

On day 10, after injecting L-DOPA with saline or ZNS (25, 50, and 100 mg/kg) for nine consecutive days (Fig 1B), AIMs were observed in different body parts and were scored for 1 min at every 20 min after L-DOPA with saline or ZNS treatment (Fig 3). We compared the total AIMs scores between chronic saline and ZNS (25, 50, and 100 mg/kg) treatments (Fig 3A; two-way repeated measures ANOVA: time, F (8, 312) = 57.44, *p* < 0.01; treatment, F (3, 39) = 1.12, *p* = 0.35; interaction, F (24, 312) = 4.80, *p* < 0.01). Chronic ZNS (50 and 100 mg/kg) treatment showed significantly higher AIMs scores than chronic saline treatment at 140 min (Tukey’s post hoc test: saline vs. ZNS (50 and 100 mg/kg), *p* < 0.01). Chronic ZNS (100 mg/kg) treatment also resulted in significantly higher AIMs scores than the other three groups at 160 and 180 min (saline vs. ZNS (100 mg/kg), *p* < 0.01; ZNS (25 and 50 mg/kg) vs. ZNS (100 mg/kg), *p* < 0.05).

**Fig 3.**
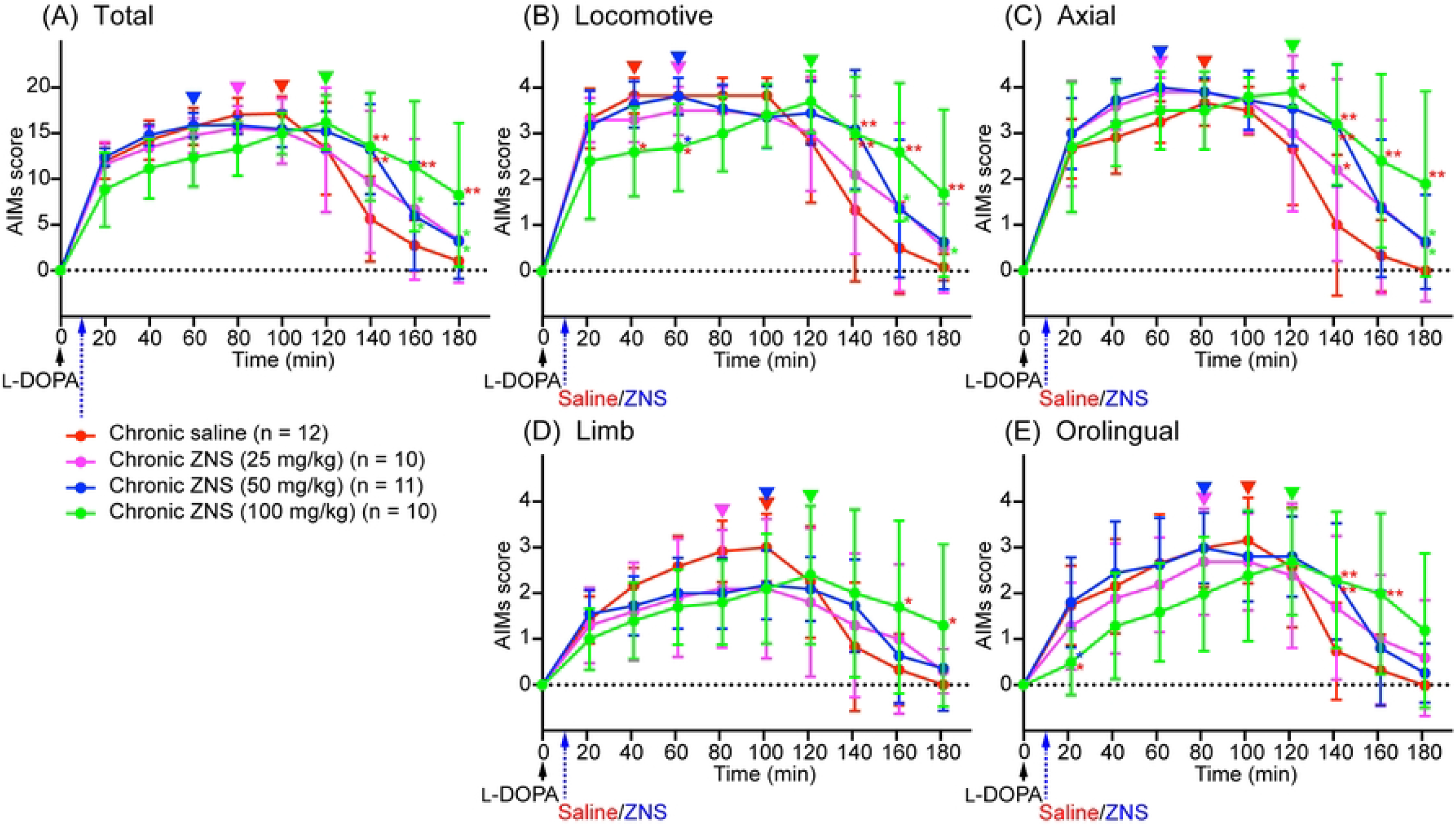
Chronic ZNS treatment delayed the peak of AIMs and decreased AIM scores before their peak, but increased their duration. Total (A), locomotive (B), axial (C), limb (D), and orolingual (E) AIMs were scored for 180 min at every 20 min after injecting L-DOPA (20 mg/kg; black arrow) at time 0 min with saline (red) or ZNS (25 mg/kg, magenta; 50 mg/kg, blue; 100 mg/kg, green) at 10 min. Data are expressed as mean ± SD. \**p* < 0.05, \*\**p* < 0.01 indicates significant differences between different treatments with the corresponding color (two-way repeated measures ANOVA followed by Turkey’s post hoc test). Peaks of AIMs with saline or ZNS are indicated by inverted triangles with the corresponding color. ZNS, zonisamide; AIMs, abnormal involuntary movements

Next, we compared the AIMs scores for each body part. In locomotive AIMs scores (Fig 3B; two-way repeated measures ANOVA: time, F (8, 312) = 49.04, *p* < 0.01; treatment, F (3, 39) = 1.01, *p* = 0.40; interaction, F (24, 312) = 4.32, *p* < 0.01), chronic ZNS (100 mg/kg) treatment showed lower AIMs scores at 40 min (Tukey’s post hoc test: saline vs. ZNS [100 mg/kg], *p* < 0.05) and 60 min (saline and ZNS [50 mg/kg] vs. ZNS [100 mg/kg], *p* < 0.05). On the other hand, chronic ZNS (100 mg/kg) treatment showed higher AIMs scores at 140 min (saline vs. ZNS [50 and 100 mg/kg], *p* < 0.01), 160 min (saline vs. ZNS [100 mg/kg], *p* < 0.01; ZNS [25 and 50 mg/kg] vs. ZNS [100 mg/kg], *p* < 0.05), and 180 min (saline vs. ZNS [100 mg/kg], *p* < 0.01; ZNS [25 mg/kg] vs. ZNS [100 mg/kg], *p* < 0.05). In axial AIMs scores (Fig 3C; two-way repeated measures ANOVA: time, F (8, 312) = 50.75, *p* < 0.01; treatment, F (3, 39) = 7.02, *p* < 0.01; interaction, F (24, 312) = 2.27, *p* < 0.01), chronic ZNS (100 mg/kg) treatment showed higher AIMs scores at 120 min (Tukey’s post hoc test: saline vs. ZNS [100 mg/kg], *p* < 0.05), 140 min (saline vs. ZNS [25 mg/kg], *p* < 0.05; saline vs. ZNS [50 and 100 mg/kg], *p* < 0.01), 160 min (saline vs. ZNS [100 mg/kg], *p* < 0.01), and 180 min (saline vs. ZNS [100 mg/kg], *p* < 0.01; ZNS [25 and 50 mg/kg] vs. ZNS [100 mg/kg), *p* < 0.05). In limb AIMs scores (Fig 3D; two-way repeated measures ANOVA: time, F (8, 312) = 31.18, *p* < 0.01; treatment, F (3, 39) = 0.19, *p* = 0.90; interaction, F (24, 312) = 4.03, *p* < 0.01), chronic ZNS (100 mg/kg) treatment showed higher AIMs scores at 160 and 180 min (Tukey’s post hoc test: saline vs. ZNS [100 mg/kg], *p* < 0.05). In orolingual AIMs scores (Fig 3E; two-way repeated measures ANOVA: time, F (8, 312) = 32.01, *p* < 0.01; treatment, F (3, 39) = 0.42, *p* = 0.74; interaction, F (24, 312) = 3.70, *p* < 0.01), chronic ZNS (100 mg/kg) treatment showed lower AIMs scores at 20 min (Tukey’s post hoc test: saline and ZNS [50 mg/kg] vs. ZNS [100 mg/kg], *p* < 0.05). In contrast, chronic ZNS (100 mg/kg) treatment resulted in higher AIMs scores at 140 min (Tukey’s post hoc test: saline vs. ZNS [50 and 100 mg/kg], *p* < 0.01) and 160 min (saline vs. ZNS [100 mg/kg], *p* < 0.01).

We compared the peaks of AIMs in the chronic saline and ZNS (25, 50, and 100 mg/kg) treatment groups (inverted triangles in Fig 3). Peaks of the total, locomotive, axial, limb, and orolingual AIMs of chronic saline and ZNS (25 and 50 mg/kg) treatments were distributed between 40 and 100 min, however those of the chronic ZNS (100 mg/kg) treatments were at 120 min.

These results indicate that chronic ZNS injection with L-DOPA in PD model mice decreased locomotive and orolingual AIMs scores before the peak and delayed the peak of the total, locomotive, axial, limb, and orolingual AIMs, but increased the duration of total, locomotive, axial, limb, and orolingual AIMs in a ZNS dose-dependent manner compared with PD model mice that underwent L-DOPA and chronic saline treatments.

## Discussion

The present study yielded the following results: (1) acute ZNS injection had no effects on L -DOPA-induced AIMs, and (2) chronic ZNS injection delayed the peak of all AIMs and decreased some AIMs scores before the peak, but increased the duration of all AIMs in a ZNS dose-dependent manner.

On day 10 after injecting L-DOPA for nine consecutive days, acute ZNS injection at all doses (25 - 100 mg/kg) did not enhance nor prolong AIMs in PD model mice (Fig 2). This result suggests that acute ZNS injection does not improve nor worsen the symptoms of LID and is consistent with a previous study on PD model rats that received L-DOPA (6 mg/kg) with ZNS (50 mg/kg) [13]. This study suggested that there was no significant difference between the L-DOPA-only and L-DOPA with ZNS groups in AIMs on day 1. A single ZNS administration may not affect LID although acute ZNS administration (300 mg) improves symptoms in PD patients [5],.

In contrast, in the present study, chronic ZNS injection (100 mg/kg) with L-DOPA delayed the peak of AIMs and decreased AIMs scores before the peak, but prolonged the duration of AIMs in a dose-dependent manner (Fig 3). Previous reports showed inconsistent effects of ZNS on AIMs in PD model rats. AIMs were observed on the 15^th^ day of L-DOPA (6 mg/kg) with ZNS (50 mg/kg) treatment after 14 consecutive injections once a day [13]. The AIMs of L-DOPA in the ZNS group were enhanced and prolonged, and the peaked AIMs score worsened slightly. On the other hand, recent reports have shown that ZNS ameliorates LID in PD model rats [18, 19], which included L-DOPA (12 mg/kg) and ZNS (26 mg/kg) injections twice a day for two weeks and AIMs evaluation. The AIMs of the L-DOPA with ZNS-injected group were statistically lower than those of the L-DOPA-only injected group 140 min after injection [18]. The effect of ZNS might be increased by the twice-daily injection of ZNS compared to the once-daily injection.

Although the precise mechanism, by which ZNS affects LID is not fully elucidated, previous studies have demonstrated that L-DOPA with ZNS treatment upregulates expression of several key molecules in the striatum of 6-OHDA-treated mice, including prodynorphin, D1 and D2 receptors, brain-derived neurotrophic factor, serotonin transporter, and 5HT1A receptor [18, 19]. Additionally, recording of neuronal activity in the SNr, one of output nuclei of the basal ganglia, in PD model mice treated with L-DOPA and ZNS injections has revealed changes in response to motor cortical stimulation [16]. In normal mice, this stimulation induces a triphasic response composed of early excitation, inhibition, and late excitation in the SNr [16, 17, 20, 21]. However, in LID model mice, longer inhibition and reduced late excitation were observed in the SNr [17]. L-DOPA with ZNS treatment further enhances these changes, which are mediated by the cortico-striato-SNr *direct* and cortico-striato-external pallido-subthalamo-SNr *indirect* pathways, respectively [20-22]. Signals through the *direct* pathway inhibit the SNr activity and release appropriate movements, while those through the *indirect* pathway excite the SNr activity and induce clear termination of movements [23, 24]. Therefore, L-DOPA with ZNS treatment enhances LID responses in the SNr and likely causes prolonged LID. ZNS increased the expression of prodynorphin and D1 receptor [18] and may enhance the activation of the *direct* pathway and cortically evoked inhibition in the SNr. ZNS inhibits low-voltage-activated Ca^2+^ currents in subthalamic nucleus neurons [25], and may reduce activity through the *indirect* pathway and cortically-evoked late excitation in the SNr.

The present study suggests that ZNS can improve LID by delaying its peak and reducing its severity before the peak although ZNS increases its duration. Additionally, ZNS may enhance the effects of L-DOPA and reduce its dosages [5-8], thereby prolonging the “honeymoon” period and delaying the appearance of LID. Further research should be investigated the optimal dose of ZNS for clinical applications.

## Acknowledgments

We thank S. Sato, K. Miyamoto, N. Suzuki, T. Sugiyama K. Awamura, and R. Kageyama for excellent technical assistance.

## References

1. Blesa J, Foffani G, Dehay B, Bezard E, Obeso JA. Motor and non-motor circuit disturbances in early Parkinson disease: which happens first? Nat Rev Neurosci. 2022;23(2):115–28. Epub 2021/12/16. doi: 10.1038/s41583-021-00542-9. PubMed PMID: 34907352.

2. Fahn S. Description of Parkinson’s disease as a clinical syndrome. Ann N Y Acad Sci. 2003;991:1–14. Epub 2003/07/09. doi: 10.1111/j.1749-6632.2003.tb07458.x. PubMed PMID: 12846969.

3. Jenner P. Molecular mechanisms of L-DOPA-induced dyskinesia. Nat Rev Neurosci. 2008;9(9):665–77. Epub 2008/08/21. doi: 10.1038/nrn2471. PubMed PMID: 18714325.

4. Kwan SY, Chuang YC, Huang CW, Chen TC, Jou SB, Dash A. Zonisamide: Review of Recent Clinical Evidence for Treatment of Epilepsy. CNS Neurosci Ther. 2015;21(9):683–91. Epub 2015/07/25. doi: 10.1111/cns.12418. PubMed PMID: 26205514; PubMed Central PMCID: PMCPMC6493109.

5. Murata M, Horiuchi E, Kanazawa I. Zonisamide has beneficial effects on Parkinson’s disease patients. Neurosci Res. 2001;41(4):397–9. Epub 2002/01/05. doi: S016801020100298X [pii]. PubMed PMID: 11755227.

6. Murata M, Hasegawa K, Kanazawa I. Zonisamide improves motor function in Parkinson disease: a randomized, double-blind study. Neurology. 2007;68(1):45–50. Epub 2007/01/04. doi: 68/1/45 [pii] 10.1212/01.wnl.0000250236.75053.16. PubMed PMID: 17200492.

7. Murata M. Novel therapeutic effects of the anti-convulsant, zonisamide, on Parkinson’s disease. Curr Pharm Des. 2004;10(6):687–93. Epub 2004/02/18. PubMed PMID: 14965331.

8. Murata M, Hasegawa K, Kanazawa I, Fukasaka J, Kochi K, Shimazu R, et al. Zonisamide improves wearing-off in Parkinson’s disease: A randomized, double-blind study. Mov Disord. 2015;30(10):1343–50. Epub 2015/06/23. doi: 10.1002/mds.26286. PubMed PMID: 26094993.

9. Binda C, Aldeco M, Mattevi A, Edmondson DE. Interactions of monoamine oxidases with the antiepileptic drug zonisamide: specificity of inhibition and structure of the human monoamine oxidase B complex. J Med Chem. 2011;54(3):909–12. Epub 2010/12/24. doi: 10.1021/jm101359c. PubMed PMID: 21175212; PubMed Central PMCID: PMCPMC3071873.

10. Sonsalla PK, Wong LY, Winnik B, Buckley B. The antiepileptic drug zonisamide inhibits MAO-B and attenuates MPTP toxicity in mice: clinical relevance. Exp Neurol. 2010;221(2):329–34. Epub 2009/12/02. doi: 10.1016/j.expneurol.2009.11.018. PubMed PMID: 19948168; PubMed Central PMCID: PMCPMC2812670.

11. Suzuki S, Kawakami K, Nishimura S, Watanabe Y, Yagi K, Seino M, et al. Zonisamide blocks T-type calcium channel in cultured neurons of rat cerebral cortex. Epilepsy Res. 1992;12(1):21–7. Epub 1992/06/01. doi: 10.1016/0920-1211(92)90087-a. PubMed PMID: 1326433.

12. Kito M, Maehara M, Watanabe K. Mechanisms of T-type calcium channel blockade by zonisamide. Seizure. 1996;5(2):115–9. Epub 1996/06/01. doi: 10.1016/s1059-1311(96)80104-x. PubMed PMID: 8795126.

13. Nishijima H, Miki Y, Ueno S, Tomiyama M. Zonisamide Enhances Motor Effects of Levodopa, Not of Apomorphine, in a Rat Model of Parkinson’s Disease. Parkinsons Dis. 2018;2018:8626783. Epub 2019/01/22. doi: 10.1155/2018/8626783. PubMed PMID: 30662707; PubMed Central PMCID: PMCPMC6312621.

14. Asanuma M, Miyazaki I, Diaz-Corrales FJ, Kimoto N, Kikkawa Y, Takeshima M, et al. Neuroprotective effects of zonisamide target astrocyte. Ann Neurol. 2010;67(2):239–49. Epub 2010/03/13. doi: 10.1002/ana.21885. PubMed PMID: 20225289.

15. Sano H, Murata M, Nambu A. Zonisamide reduces nigrostriatal dopaminergic neurodegeneration in a mouse genetic model of Parkinson’s disease. J Neurochem. 2015;134(2):371–81. Epub 2015/04/11. doi: 10.1111/jnc.13116. PubMed PMID: 25857446.

16. Sano H, Nambu A. The effects of zonisamide on L-DOPA-induced dyskinesia in Parkinson’s disease model mice. Neurochem Int. 2019;124:171–80. Epub 2019/01/15. doi: 10.1016/j.neuint.2019.01.011. PubMed PMID: 30639196.

17. Dwi Wahyu I, Chiken S, Hasegawa T, Sano H, Nambu A. Abnormal Cortico-Basal Ganglia Neurotransmission in a Mouse Model of l-DOPA-Induced Dyskinesia. J Neurosci. 2021;41(12):2668–83. Epub 2021/02/11. doi: 10.1523/JNEUROSCI.0267-20.2020. PubMed PMID: 33563724; PubMed Central PMCID: PMCPMC8018735.

18. Oki M, Kaneko S, Morise S, Takenouchi N, Hashizume T, Tsuge A, et al. Zonisamide ameliorates levodopa-induced dyskinesia and reduces expression of striatal genes in Parkinson model rats. Neurosci Res. 2017;122:45–50. Epub 2017/06/05. doi: 10.1016/j.neures.2017.04.003. PubMed PMID: 28577977.

19. Tohge R, Kaneko S, Morise S, Oki M, Takenouchi N, Murakami A, et al. Zonisamide attenuates the severity of levodopa-induced dyskinesia via modulation of the striatal serotonergic system in a rat model of Parkinson’s disease. Neuropharmacology. 2021;198:108771. Epub 2021/09/03. doi: 10.1016/j.neuropharm.2021.108771. PubMed PMID: 34474045.

20. Chiken S, Sato A, Ohta C, Kurokawa M, Arai S, Maeshima J, et al. Dopamine D1 Receptor-Mediated Transmission Maintains Information Flow Through the Cortico-Striato-Entopeduncular Direct Pathway to Release Movements. Cereb Cortex. 2015;25(12):4885–97. Epub 2015/10/08. doi: 10.1093/cercor/bhv209. PubMed PMID: 26443442; PubMed Central PMCID: PMCPMC4635926.

21. Sano H, Chiken S, Hikida T, Kobayashi K, Nambu A. Signals through the striatopallidal indirect pathway stop movements by phasic excitation in the substantia nigra. J Neurosci. 2013;33(17):7583–94. Epub 2013/04/26. doi: 10.1523/JNEUROSCI.4932-12.2013. PubMed PMID: 23616563; PubMed Central PMCID: PMCPMC6619573.

22. Tachibana Y, Kita H, Chiken S, Takada M, Nambu A. Motor cortical control of internal pallidal activity through glutamatergic and GABAergic inputs in awake monkeys. Eur J Neurosci. 2008;27(1):238–53. Epub 2007/12/21. doi: 10.1111/j.1460-9568.2007.05990.x. PubMed PMID: 18093168.

23. Nambu A, Tokuno H, Takada M. Functional significance of the cortico-subthalamo-pallidal ’hyperdirect’ pathway. Neurosci Res. 2002;43(2):111–7. Epub 2002/06/18. doi: 10.1016/s0168-0102(02)00027-5. PubMed PMID: 12067746.

24. Mink JW, Thach WT. Basal ganglia intrinsic circuits and their role in behavior. Curr Opin Neurobiol. 1993;3(6):950–7. Epub 1993/12/01. doi: 10.1016/0959-4388(93)90167-w. PubMed PMID: 8124079.

25. Yang YC, Tai CH, Pan MK, Kuo CC. The T-type calcium channel as a new therapeutic target for Parkinson’s disease. Pflugers Arch. 2014;466(4):747–55. Epub 2014/02/18. doi: 10.1007/s00424-014-1466-6. PubMed PMID: 24531801.

